# Transcriptome analysis of genetically matched human induced pluripotent stem cells disomic or trisomic for chromosome 21

**DOI:** 10.1101/100859

**Authors:** Patrick Gonzales, Gin Fonte, Christine Roberts, Connor Jacobsen, Gretchen H. Stein, Christopher D. Link

**Author notes:** Corresponding author: Christopher D. Link, Ph.D., Associate Professor, Linda Crnic Institute/Integrative Physiology, University of Colorado, Campus Box 354 Boulder, CO 80309.

## Abstract

Trisomy of chromosome 21, the genetic cause of Down syndrome, has the potential to alter expression of genes on chromosome 21, as well as other locations throughout the genome. These transcriptome changes are likely to underlie the Down syndrome clinical phenotypes. We have employed RNA-seq to undertake an in-depth analysis of transcriptome changes resulting from trisomy of chromosome 21, using induced pluripotent stem cells (iPSCs) derived from a single individual with Down syndrome. These cells were originally derived by Li et al, who genetically targeted chromosome 21 in trisomic iPSCs, allowing selection of disomic sibling iPSC clones. Analyses were conducted on trisomic/disomic cell pairs maintained as iPSCs or differentiated into cortical neuronal cultures. In addition to characterization of gene expression levels, we have also investigated patterns of RNA adenosine-to-inosine editing, alternative splicing, and repetitive element expression, aspects of the transcriptome that have not been significantly characterized in the context of Down syndrome. We identified significant changes in transcript accumulation associated with chromosome 21 trisomy, as well as changes in alternative splicing and repetitive element transcripts. Unexpectedly, the trisomic iPSCs we characterized expressed higher levels of neuronal transcripts than control disomic iPSCs, and readily differentiated into cortical neurons, in contrast to another reported study. Comparison of our transcriptome data with similar studies of trisomic iPSCs suggests that trisomy of chromosome 21 may not intrinsically limit neuronal differentiation, but instead may interfere with the maintenance of pluripotency.

## Introduction

Down Syndrome (DS) results from an extra copy of chromosome 21, and this change in gene dosage has been proposed to alter chromosome 21 gene expression. Chromosome 21 trisomy also has the potential to alter the global transcriptome, either by secondary effects of chromosome 21 gene over-expression, or as a byproduct of additional genetic material itself. In addition to the possible perturbation of specific cellular pathways by altered expression of chromosome 21 genes, chromosome 21 also contains genes that impact the global transcriptome directly. These include *ADARB1,* encoding one of two genes responsible for adenosine-to-inosine RNA editing; U2AF1, a constitutive splicing factor; and DYRK1A, a kinase known to target splicing factors. Identifying transcriptome changes reproducibly caused by chromosome 21 trisomy may be crucial to understanding the origins of the clinical features of Down Syndrome.

Initial gene expression studies that sought to establish chromosome 21-dependent transcriptome changes using human material were confounded by the effects of different genetic backgrounds in the DS and control individuals who were studied (Gardiner et al, 2010). An alternative approach is to characterize gene expression in inbred mouse DS models, which reproduce some of the developmental and behavioral phenotypes of DS individuals (Liu et al, 2011). Transcriptome studies have been done in multiple mouse DS models (Saran et al, 2003; Potier et al, 2006; Ling et al, 2014; Guedj et al, 2015; Olmos-Serrano et al, 2016), but no consistent pattern of transcriptome changes has been identified, perhaps due to the different tissues and developmental stages analyzed in these studies. In addition, these mouse DS models have triplication of chromosomal regions syntenic to only part of human chromosome 21 as well as triplication of non-chromosome 21 homologous genes, and thus may not capture the full effects of human chromosome 21 trisomy.

Recently, the development of paired disomic/trisomic induced pluripotent stem cells (iPSCs) derived from the same trisomic individual (or from discordant monozygotic twins) has allowed the measurement of human trisomy 21-dependent gene expression independent of genetic background differences (Li et al, 2012: Weick et al, 2013; Jiang et al, 2013; LeTourneau et al, 2014). These studies have consistently observed a general upregulation of genes on chromosome 21, as well as highly variable significant gene expression differences on other chromosomes. Li et al (2012) used an elegant genetic selection system to derive disomic iPSC subclones from chr21 trisomic iPSCs that were derived from an individual with Down syndrome. Based on an initial microarray analysis, these researchers also reported a general upregulation of chromosome 21 genes, but could identify only 3 non-chromosome 21 genes with a > 2 fold dysregulation between disomic and trisomic iPSC pairs. We have used RNA-seq to significantly extend this analysis using both disomic/trisomic iPSC pairs and cortical neuron cultures derived from them. In addition to assaying transcript accumulation, we have used the RNA-seq data to look for other possible trisomy-induced transcriptome changes, including possible alterations in adenosine-to-inosine RNA editing, alternative splicing, and repetitive element expression.

In contrast to the Jiang et al study, the trisomic iPSCs we characterized were not restricted in their ability differentiate into cortical neuron cultures, and in fact showed higher expression of neuronal transcripts when compared to disomic iPSCs. To investigate differences between our analyses and previous studies, we have compared global gene expression data among the related studies, and find evidence for at least two different states for trisomic iPSC. We suggest that these different states may reflect an inherent inability for trisomic iPSCs to maintain full pluripotency.

## Materials and Methods

### iPSC Cell Culture

Trisomic (C2 and C3) and disomic iPSC cell lines (C2-4-3, C2-4-4, C3-5-11, and C3-5-13), which had been developed from an individual with Down syndrome by Li et al. (2012), were obtained through the Linda Crnic Institute for Down Syndrome (University of Colorado, Aurora, CO). These iPSC lines were grown on Geltrex (Life Technologies) under feeder-free conditions in mTeSR1 medium according to the protocol of Stem Cell Technologies. iPSC colonies were passaged with dispase, and differentiating colonies were removed manually by scraping. Periodically, live iPSC colonies were stained with anti-Tra-1-60 or Tra-1-81 conjugated to Dylight fluor 488 to confirm the stem cell character of the colonies. Likewise, fixed cells were shown to express Oct4 by immunostaining. We used ABI/Thermo Fisher TaqMan Copy Number Assays (Hs01180853_cn; Hs02928366_cn; Hs01533676 corresponding to APP, DYRK1A and RCAN1 respectively) to evaluate the number of copies of Chr. 21 in the iPSC lines. We found that C2 was stably trisomic, but that C3 appeared initially to be a mixture of trisomic and disomic cells that rapidly gave rise to a consistently disomic cell line, which we call C3-D21.

### Neuronal Differentiation

iPSC cells were induced to differentiate to cortical neurons by a modification of the method of Espuny-Camacho et al. (2013). Briefly, iPSCs were dissociated with Accutase, and plated on Geltrex-coated dishes at 10-20 × 10^3^ cells/cm^2^ in mTeSR1 medium + 10 uM Y27632 ROCK inhibitor (Tocris). After 2-4 days, the cells formed a meshwork of dense ridges with mostly open spaces in between. The differentiation process was begun (day 0) by feeding the cells every other day for 12 days with NIM neural induction medium (Stem Cell Technologies), followed by DDM medium supplemented with 2% B27 (Gaspard et al., 2009) every other day for 12 days. On day 24, the dense ridges or islands containing neural progenitors were manually dissociated, resuspended in DDM + 2% B27 + 10 uM Y27632, and plated on dishes and chamber slides coated with poly-ornithine and laminin (Shi et al., 2012). Every 4-5 days, half the medium was changed to fresh Neurobasal + 2% B27 + 2mM GlutaMax (Invitrogen). Colonies that were clearly nonneuronal were removed by scraping, rinsing and refeeding the cultures. After 40 days of differentiation, the cells were rinsed with PBS and harvested with ice-cold Trizol or fixed with 4% para-formaldehyde + 4% sucrose in PBS.

Successful differentiation of iPSCs into cortical neuronal cultures was confirmed by immunofluorescence staining for markers of cortical layers V (transcription factor Ctip2, encoded by BCL11B) and VI (transcription factor TBR1), as well as for glutamatergic markers vGLUT1 (encoded by SLC17A7) and GRIK2 (see Supplemental Figure 1).

### Immunofluorescence staining

For live cell staining, StemGent antibodies conjugated to Dylight 488: Tra1-60 or Tra1-81, were applied to iPSC cultures for 30-60 min according the manufacturer’s instructions, and the stained colonies were examined with an Evos FL microscope.

For staining of fixed cells, cells were grown on chamber slides (LabTek II CC2), rinsed with Dulbecco’s PBS, fixed 15 min in 4% p-formaldehyde/4% sucrose in PBS, permeabilized 7 min in 0.25% Triton X-100 in PBS, and blocked 1-2 hrs with 5% sheep or goat serum in PBS. Primary antibodies, applied overnight at 4°C, were from Abcam: Oct4 (ab19857), Tbr1 (ab31940), Ctip2 (ab18465),GRIK2 (ab53092), vGlut1 (ab72311) and βIII-tubulin (ab7751); from Millipore: MAP2 (polyclonal AB5622), MAP2 (monoclonal MAB3418). The slides were rinsed (5X PBS), secondary antibodies (Life Technologies) were applied for 2 hrs at room temperature, rinsed (4X PBS), stained with DAPI (Sigma D9542), and mounted with ProLong Gold AntiFade (Life Technologies). Images were captured using an epifluorescence microscope (Zeiss Axioskop) equipped with a CCD camera and Slidebook image analysis software (3i).

### Immunoblotting

Lysates of disomic and trisomic iPSCs were prepared with RIPA buffer containing protease and phosphatase inhibitor cocktails (Sigma), and the protein content of each lysate was determined (Bradford assay). Using BioRad reagents and equipment, an equal amount of protein from each lysate was denatured by boiling 5 minutes in 4x Sample Buffer containing DTT; run on a Criterion XT pre-cast gradient gel (4%-12%) in XT-MOPS running buffer at 150 volts for 1.5-2 hours; and blotted via a Trans-Blot Turbo system for 7 minutes using the Mixed Molecular Weight protocol (2.5A, up to 25V). The blot was washed in TBS-T, (tris-buffered saline containing Tween 20), stained with Ponceau S to affirm protein transfer, washed again and blocked in 5% milk in TBS-T before being stained sequentially with MAP2 rabbit polyclonal antibody (1:1000, Abcam ab32454) and goat anti-rabbit HRP (1:1000). The washed blot was then treated with SuperSignal West Femto Maximum Sensitivity substrate (Life Technologies), diluted 1:10 in distilled water, for 5 minutes while covered, and images were taken using a ChemiDoc-it imaging system and VisionWorks LS software. The blot was stripped and stained for GAPDH (1:2000) as a loading control. Images were analyzed using Fiji (ImageJ) and Excel; integrated density of each band was measured and normalized to the GAPDH signal before comparing the amount of MAP2 in each protein sample.

### RNA isolation, cDNA library preparation, high-throughput sequencing

RNA for poly(A) RNA sequencing libraries was extracted from IPSC and neuronal cultures via TRIzol extraction. Genomic DNA was removed using TURBO DNase (Invitrogen). RNA integrity (RIN) was verified on an Agilent 2100 bioanalyzer and only samples with RIN ≥ 7.0 were used for sequencing. Single-end sequencing libraries were prepared by the UCCC Genomics Core (Aurora, CO) using the TruSeq Stranded mRNA Library Prep Kit (Illumina). Cluster generation and sequencing were performed on the Illumina HiSeq 2000 platform. The reads were de-multiplexed and converted to FASTQ format using CASAVA software from Illumina.

### Analysis of RNA-seq data

Analysis of library complexity and per-base sequence quality (i.e. q > 30) was assessed using FastQC (v0.11.2) software (Andrews, 2010). Low quality bases (q < 10) were trimmed from the 3’ end of reads. Adaptor sequences and reads shorter than 40 nucleotides were removed using Trimmomatic (Bolger, 2014). Reads were aligned to GRCh37/hg19 using TopHat2 (v2.0.14, --b2-very-sensitive --keep- --no-coverage-search --library-type fr-firststrand) (Kim et al.,2013). High quality mapped reads (MAPQ > 10) were filtered with SAMtools (v0.1.19) (Li et al., 2009). Gene level counts were obtained using the GRCh37 Ensembl annotation with Rsubread (v 1.18.0, strandSpecific = 2, GTF.featureType = “exon”, countMultiMappingReads = FALSE) (Liao, 2013). Differential expression of genes was determined using DESeq (v1.24.0) software with the options “per-condition”, “maximum”, and “local” for dispersion calculations (Anders, 2010). Significance was assigned to genes with an FDR less than 10%.

### Global RNA Editing

The bioinformatics scheme was implemented to specifically quantify the amount of RNA editing that occurs in known RNA editing locations. With this is mind we used the “DAtabase of RNa Editing” (DARNED) to procure a list of all previously published RNA editing locations in the human genome (hg19) (Kiran, 2010). Only Adenosine to Inosine RNA editing sites in exons were used in the analysis. Reads overlapping DARNED locations were kept if they were uniquely aligned with a MAPQ greater than 10, and a minimum PHRED score of 30 at the nucleotide level. An edit ratio was then determined for each location by summing the total number of edits (A to G changes) divided by the total possible edits (read depth at edit location). A minimum read depth of 10 was required for the edit ratio to be determined. The global editing percentage for each sample was then ascertained by averaging all edit ratios considered.

### Alternative splicing analysis

Splicing analysis was performed using Hartley’s QoRTs / JunctionSeq pipeline (Hartley, 2016). Tophat aligned reads and an Ensembl annotation file (GRCh37) were used as input to generate gene counts with QoRTs software using the options “--stranded”, “--singleEnded”, and “--minMAPQ 50”. A flat annotation file containing known splice junctions was produced in QoRTs using the same Ensembl annotation above with the “--stranded” option set. The QoRTs generated files were both used as input into JunctionSeq for differential exon and splice junction analysis. Exons or splice junctions were considered differentially expressed in the disomic vs. trisomic comparison if the adjusted P value was less than 10%.

### Semi-quantitative PCR

To capture the exclusion of APOO exon 4 in trisomic samples we designed primers bordering exon 3 and 5 (FWD 5’ TCTCACAGCTCCGACACTAT 3’, REV 5’ GAGTCCAATAAGGCCAGCAA 3’). 30ng/ul of cDNA were used as input for all samples. PCR cycling conditions were set as follows: denaturation for 5 min at 95 °C, followed by 40 cycles of denaturation for 30 s at 94 °C, annealing for 30 s at 54 °C, and polymerization for 30s at 72 °C, and a final extension for 10 min at 72 °C. APOO exon 4 exclusion was resolved on 2% agarose gel.

### Expression of repetitive elements

RepEnrich software was used to generate a count table containing uniquely aligned reads assigned to repetitive elements (Criscione et al., 2014). An hg19 repeatmasker file was used to build the repetitive element annotation. Size factors generated from the DESeq gene expression analysis were used to normalize the samples. DESeq (v1.24.0) was used to determine differential expression of repetitive element transcripts. Significance was assigned to genes with an FDR less than 10%.

### Cluster analysis of datasets

The full datasets of normalized transcript accumulation values for the iPSC studies by Weick et al, 2013; Jiang et al, 2013; and LeTourneau et al, 2014 were downloaded from the Gene Expression Omnibus (GSE48611, GSE47014, and GSE55504, respectively). Trisomic/disomic ratios were calculated using the averaged transcript levels for all genes assayed in each study. Ratios were imported into Cluster 3.0 (de Hoon et al, 2004) for genes identified as significantly differentially expressed in our study, and present in 6/7 of the studies being compared. The ratios were log-transformed, and hierarchical clustering was performed using centroid linkage and centered correlation for the similarity matrix. The clustering output was visualized using Java TreeView (Saldanha, 2004).

## Results

### Generation of RNA-seq data

Paired trisomic/disomic iPSCs were obtained from the Russell lab and maintained in feeder-free media. Cortical neuronal cultures were derived from the iPSCs following the procedure of Espuny-Camacho et al (2013) with minor modifications (see Materials and Methods) and harvested after 40 days in culture. All sequence data was generated from polyA-selected, strand-specific sequencing libraries by Illumina sequencing (100-115 bp single end reads). The iPSC data was generated from 3 independent preps of trisomic clone C2 from the Russell lab, and 3 independent preps of disomic sub-clones derived from the C2 lineage (two C2-4-4 and one C2-4-3). Our analysis of the transcriptomes of cortical neurons compared 3 biologically independent sets of differentiated cortical neurons derived from the trisomic C2 line and from disomic C3 iPSC cells (C3-D21, which arose spontaneously from trisomic clone C3). Differentially expressed transcripts were identified using the DESeq algorithm. Principal component analysis demonstrated that the 12 transcriptome datasets partitioned as expected (see Figure 1).

**Figure 1.**
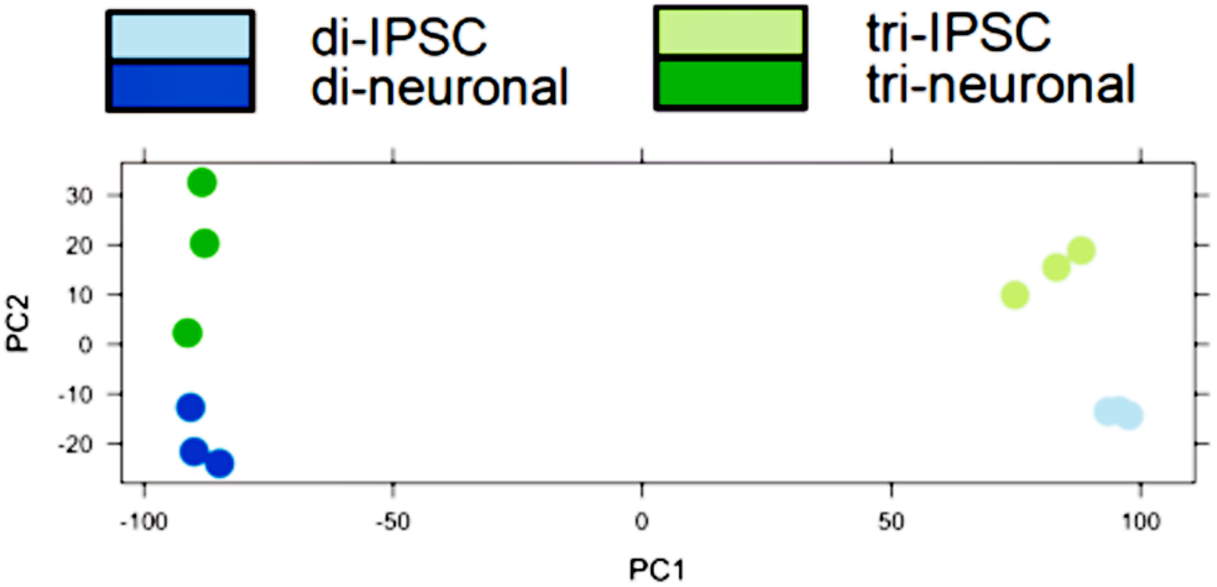
Principal component analysis of transcriptome datasets. Note principal component 1 clearly segregates iPSC from neuronal cultures, while component 2 partitions trisomic from disomic cells.

### Gene expression in trisomic cells

Using a false discovery rate (FDR) of 10%, we identified 1644 transcripts (810 up, 834 down) with differential expression between trisomic and disomic iPSCs (see Supplementary table 1). Gene ontology analysis revealed that trisomic iPSCs had increased transcript levels for genes involved in neurogenesis and neuronal function (see Table 1). In contrast, genes with decreased transcripts in the trisomic iPSCs were heavily over-represented in cell adhesion function and germ layer/mesoderm development. The germ layer category includes genes well-established to function in maintenance of stem cell fates (e.g., KLF4, NODAL, LEFTY1 and 2, etc).

**Table 1.**
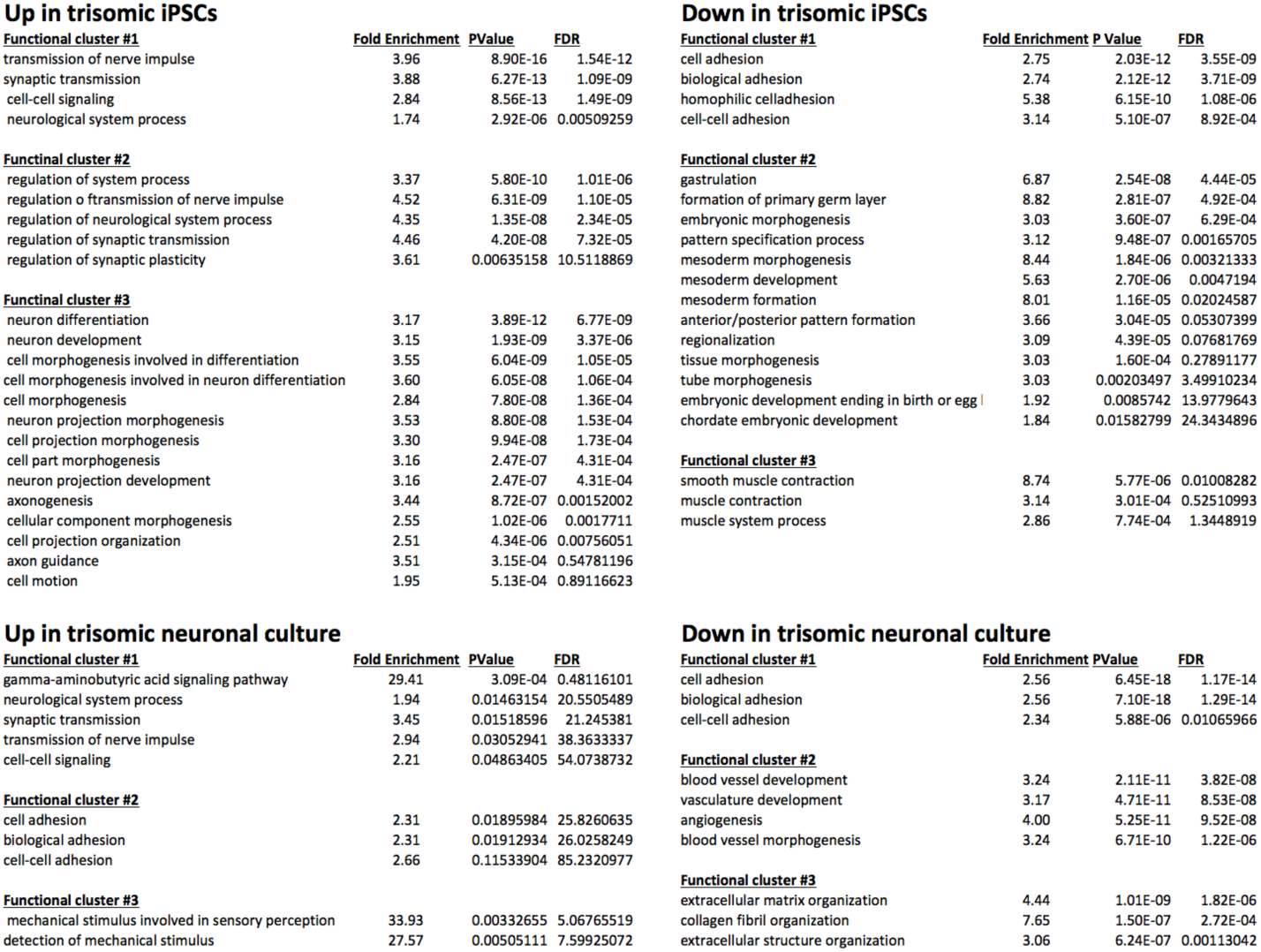
Gene ontology enrichment using DAVID *(https://d9vid.ncifcrf.gov/).*

To confirm the surprising observation that the trisomic iPSCs appeared to express neuronal transcripts at higher levels than the paired disomic iPSCs, we focused on MAP2, a classic neuronal-specific microtubule-binding protein whose transcript was increased an average of 6 fold in the trisomic iPSCs compared to the disomic iPSCs. As shown in Figure 2, MAP2 protein was also significantly increased in the trisomic iPSCs as assayed by either immunoblot or immunofluorescence.

**Figure 2.**
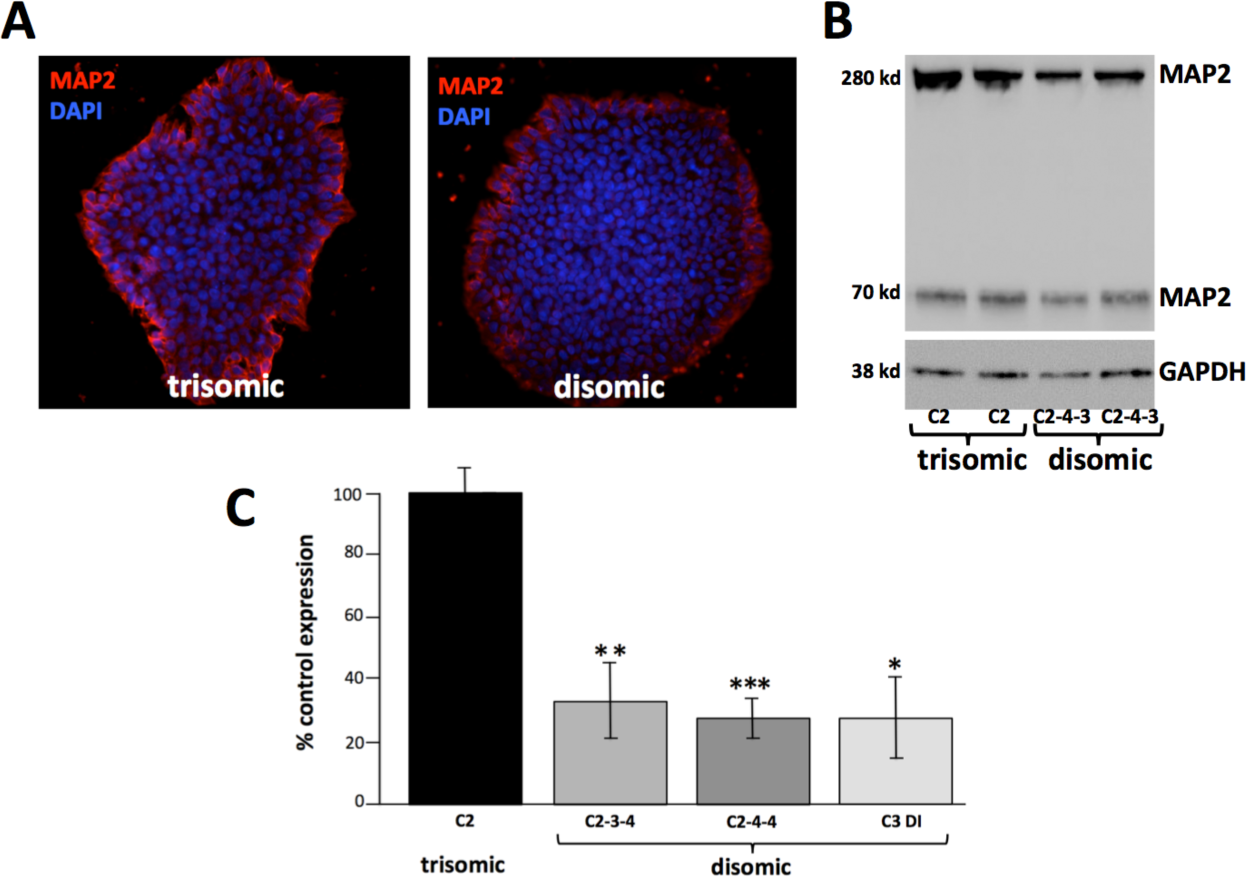
Trisomic iPSCs show higher levels of MAP2 expression. **A.** iPSC colonies fixed and stained for MAP2. **B.** Representative immunoblot from trisomic and disomic iPSCs. Note two MAP2 isoforms are detected. **C.** Quantitation of MAP2 immunoblots. (* = P<0.02, ** = P< 0.01, ***, P< 0.001, paired T-tests)

Letourneau et al (2014) claimed to identify gene expression dysregulation domains (GEDDs) in trisomic iPSCs and fibroblasts. We applied the same algorithmic approach to our iPSC data, and either did not see the same patterns of expression domains (see Supplemental Figure 2) or did not detect statistically significant expression domains.

Using the same parameters applied in the iPSC comparison to the RNA-seq data from the neuronal cultures, we identified 1,377 transcripts (212 up in trisomic cells, 1165 down) differentially expressed in the trisomic and disomic neuronal cultures (See Supplementary Table 2). Gene ontology analysis of differentially expressed transcripts (Table 1) identified similar functional categories as observed in the precursor iPSCs. Transcripts for many GABA and glutamate receptors were markedly increased in the trisomic neuronal cultures. We note that transcripts for general neuronal markers (e.g., MAP2, βIII tubulin, and SNAP25) were not increased in the trisomic neuronal cultures compared to the disomic neuronal cultures, suggesting that the increases in synaptic markers in the trisomic neuronal cultures were not simply the result of a greater proportion of neurons in these cultures, but rather reflect functional differences in the neurons themselves. This observation seems to differ from the results of Weick et al (2013), who observed reduced synaptic activity in their trisomic iPSC-derived neuronal cultures. As observed for the iPSCs, many categories of cell adhesion molecules showed decreased transcript accumulation in the trisomic neuronal cultures. (One class of cell adhesion-associated transcripts was increased in the trisomic neuronal cultures, however, this class encodes predominantly neuronal-specific adhesion molecules such as *DAB1, CDH18,* and *NELL2.)* As expected, stem cell-associated transcripts were expressed at low levels in both the trisomic and disomic neuronal cultures, with no significant differential expression.

In both the iPSC and neuronal trisomic cultures, chromosome 21 transcripts were increased overall, at close to the 1.5 fold increase predicted from a simple gene dosage effect, however, the specific chromosome 21 genes over-expressed was dependent on the cell types being analyzed (Figure 3)

**Figure 3.**
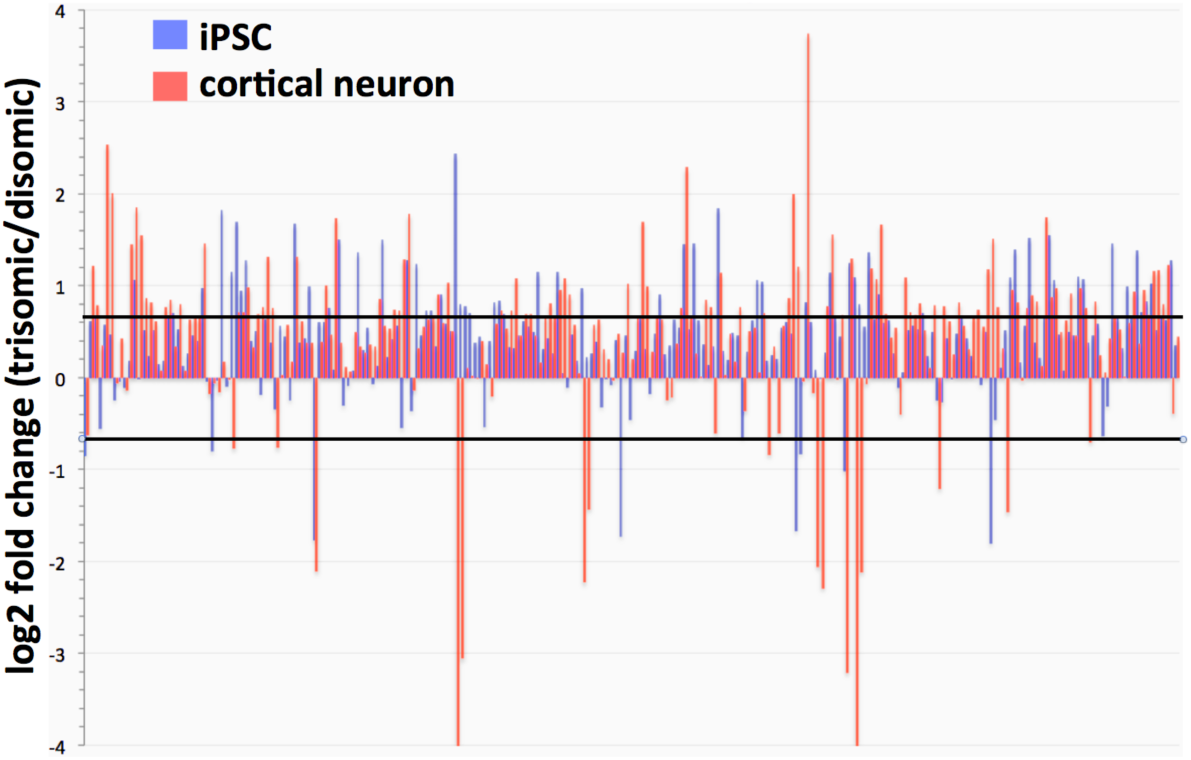
Expression of Chr 21 genes in iPSCs and cortical neuronal cultures (trisomic/disomic ratios). Lines represent individual gene ratios across chromosome 21. Note strong general trend for Chr 21 genes to be over-expressed, although specific genes may be strongly up- or-down regulated depending on cell type. (Black lines indicate 1.5 fold increases or decreases in trisomic cells.)

### Adenosine-to-inosine (A-to-I) editing in trisomic cells

ADARB1 encodes one of the two enzymes responsible for post-transcriptional A-to-I editing of RNA. This gene is encoded on chromosome 21, leading to suggestions that excessive RNA editing may occur in trisomy 21 cells. To test this hypothesis, we scored editing levels at ~10,000 residues annotated to undergo A-to-I editing (the DARNED database) in the trisomic and disomic neuronal cultures. (We did not score editing in the iPSCs, as A-to-I editing happens preferentially in neuronal cells, and in fact the iPSCs had very low levels of ADARB1 transcripts.) We found that ADARB1 transcripts are increased in the trisomic neuronal cultures, however levels of these transcripts do not correlate well with over all levels of A-to-I editing (see Figure 4). We conclude that increased levels of ADARB1 transcripts do not lead to increases in global RNA editing. Our dataset does not have sufficient read depth to determine rigorously if there were significant changes in site-specific editing in the trisomic neuronal cultures.

**Figure 4.**
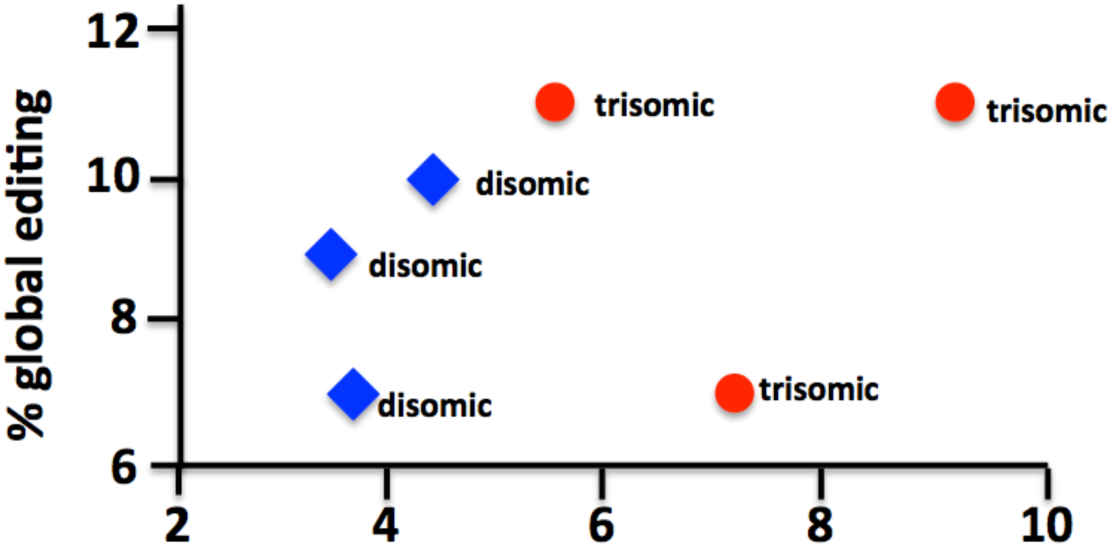
Correlation between ADARB1 transcript abundance and levels of global A-to-I editing. Note that all trisomic cultures had higher normalized levels of ADARB1 transcripts, but this did not necessarily translate into higher editing rates.

### Splicing alterations in trisomic cells

To identify differences in splicing between trisomic and disomic cells, we tested multiple splicing algorithms, and chose JunctionSeq (see Methods). Using this algorithm we identified 117 annotated genes with splicing changes when comparing trisomic and disomic iPSCs, and 36 such genes when comparing the derived cortical neuron cultures (using a conservative adjusted P value < 0.01). (See Supplemental Table 3.) Only one gene, SLC38A2, appeared to have altered splicing in both the iPSCs and cortical neuronal cultures, perhaps not surprising given the large transcriptional and splicing differences between stem cells and neurons. To verify this bioinformatic analysis, we performed semi-quantitative RT-PCR on a subset of genes identified in the iPSC analysis. Shown in Figure 5 are the results for the Apolipoprotein O gene (APOO). As predicted by the JunctionSeq analysis, trisomic cells show increased exclusion of exon 14.

**Figure 5.**
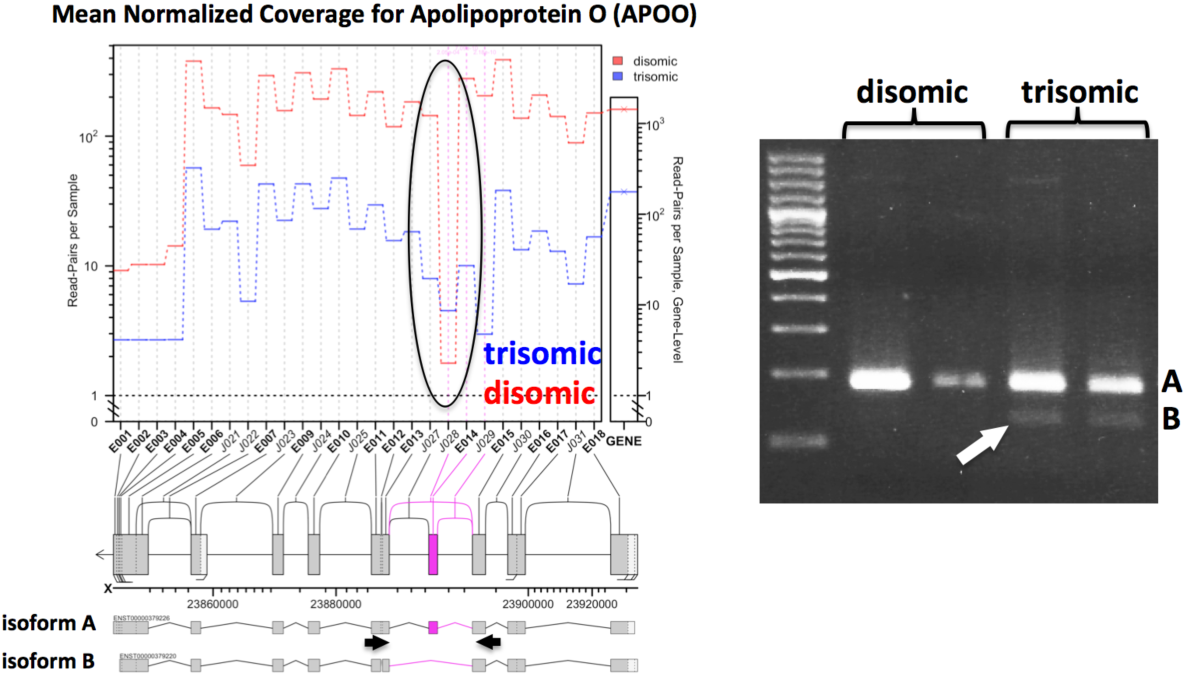
Altered splicing pattern of APOO in trisomic cells. **A.** JunctionSeq output supporting the skipping of exon 14 in trisomic iPSCs. **B.** RT-PCR targeting exon 14 exclusion (primer locations shown by arrows in panel A). Arrow indicates exon exclusion band only recovered in trisomic cells.

### Accumulation of repetitive element transcripts in trisomic cells

Retrotransposons, one class of repetitive elements, appear to play an important role in the maintenance of stem cell pluripotency (Macfarlan et al, 2012; Fort et al, 2014). To determine whether chromosome 21 trisomy might impact the accumulation of repetitive element (RE) transcripts globally, we used the RepEnrich algorithm (Criscione et al, 2014) to compare RE transcript levels in both the iPSC and differentiated neuronal trisomic/disomic culture pairs (see Supplemental table 4). In the iPSC comparison, there was a trend towards decreased RE transcript accumulation in the trisomic cells, with 26/38 differentially expressed RE transcripts (FDR <10%) lower in the trisomic cells. Interestingly, the most highly expressed RE transcript in the iPSCs was HERVH-int, which also showed the most (statistically) significant difference between the trisomic and disomic iPSCs: a > 3 fold reduction in the trisomic cells. There was no significant difference in HERVH-int expression in cortical neuronal cultures, and the overall trend in RE expression appeared reversed, with 18/21 significantly different RE transcripts increased in the trisomic cells.

### Comparison to other transcriptome data sets

The iPSCs and induced neuronal cultures that we have characterized were derived from a single individual, thereby intrinsically limiting the generalization of our results. However, three datasets have been published that have attempted a similar transcriptome characterization using trisomic/disomic iPSC pairs that have the same genetic background within each study (Table 2). Weick et al (2013) derived iPSCs from fibroblasts from a DS individual who was mosaic for chromosome 21 trisomy, which allowed the derivation of both trisomic and disomic iPSCs. Jiang et al (2013) used an elegant, inducible XIST-based system to inhibit expression of one chromosome 21 in trisomic iPSC clones, which also allowed a trisomic/disomic comparison in the same iPSC clone. Letourneau et al (2014) identified a rare set of monozygotic twins discordant for chromosome 21 trisomy, and that enabled them to derive a trisomic/disomic iPSC pair. As we have done, these studies all used statistical criteria to generate lists of genes with differential expression in trisomic vs. disomic iPSCs. Comparison of these lists identified no differentially expressed gene common to all the studies, except for genes expressed on chromosome 21. There are multiple, non-mutually exclusive, possible explanations for this observation: inherent variability in independently-generated iPSCs, large effects of the experimental conditions used in individual labs, insufficient replication to identify all transcriptome differences accurately, strong effects of genetic background, one or more study with outlying data, or a fundamental absence of a non-chromosome 21 transcriptome signature in trisomic iPSCs. To sort through these possibilities, we undertook a cluster analysis to determine how the datasets grouped. In particular, we sought to determine if our dataset was truly an outlier in comparison to the previous studies.

**Table 2.** Comparable studies assaying transcriptome differences in isogenic disomic/trisomic iPSCs pairs.

| <u>study</u> | <u>isogenic iPSC approach</u> | <u>sex</u> | <u>transforming factors</u> | <u>culture</u> | <u>transcriptome analysis</u> |
| --- | --- | --- | --- | --- | --- |
| Li et al, 2012<br>(used in our study) | genetic selection for Chr 21 loss | M | LIN28, NANOG, SOX2, OCT4 | feeder-free | RNA-seq |
| Weick et al, 2013 | fibroblasts from mosaic individual | M | OCT4, SOX2, KLF4, MYC | MEFs | microarray |
| Jiang et al, 2013 | inducible XIST inactivation of one Chr 21 | M | OCT4, SOX2, KLF4, MYC | MEFs | microarray |
| Letourneau et al, 2014 | fibroblasts from discordant twin pair | F | OCT4, SOX2, KLF4, MYC | human foreskin | RNA-seq |

The relevant data from previous studies were downloaded from online repositories, and trisomic/disomic ratios were calculated for all the annotated genes assayed. This approach enabled us to use a dimensionless measure that allowed comparison of RNA-seq and microarray data. Our study and the twin study by Letourneau et al (2014) produced single sets of trisomic/disomic ratios. The study by Weick et al (2013) generated two sets of ratios, using the data from two trisomic clones (DS1 and DS4) and one disomic clone (DS2). (We calculated the DS1/DS2 and DS4/DS2 ratios to maximize information, while appreciating that these ratios are not independent because they use the same denominator.) The study by Jiang et al (2013) examined 3 independent XIST chromosome 21 insertion subclones derived from the same parental trisomic iPSC clone. These subclones were subjected to doxycyline-induced XIST silencing, and trisomic/disomic ratios could be calculated by comparing uninduced/induced transcriptome measurements, thus generating 3 independent ratios (clones 1, 2, 3). The cluster analysis was based on the trisomic/disomic ratios of our set of differentially expressed genes and the calculated ratios for the same genes in the other datasets. After filtering for genes with values in at least 5/6 of the other datasets, 942 genes were used for the clustering analysis (see Materials and Methods).

Figure 6A shows the dendrogram of the calculated dataset relationships. Note that our dataset does not appear to be an outlier, but instead clusters with clone 3 of the study by Jiang et al (2013) and with the twin study of Letourneau et al. (2014). Figure 6B shows a portion of the gene expression heat map for the clustered datasets, capturing genes that had increased expression (red color) in our dataset (second column in heat map display). Note that these genes include neuron-associated transcripts such as neurexin (NRXN1), neurocan (NCAN), and α-2A adrenoreceptor (ADRA2A), all of which also show increased expression in the trisomic iPSCs characterized in the twin study of Letourneau et al (2014; first column in heat map display).

**Figure 6.**
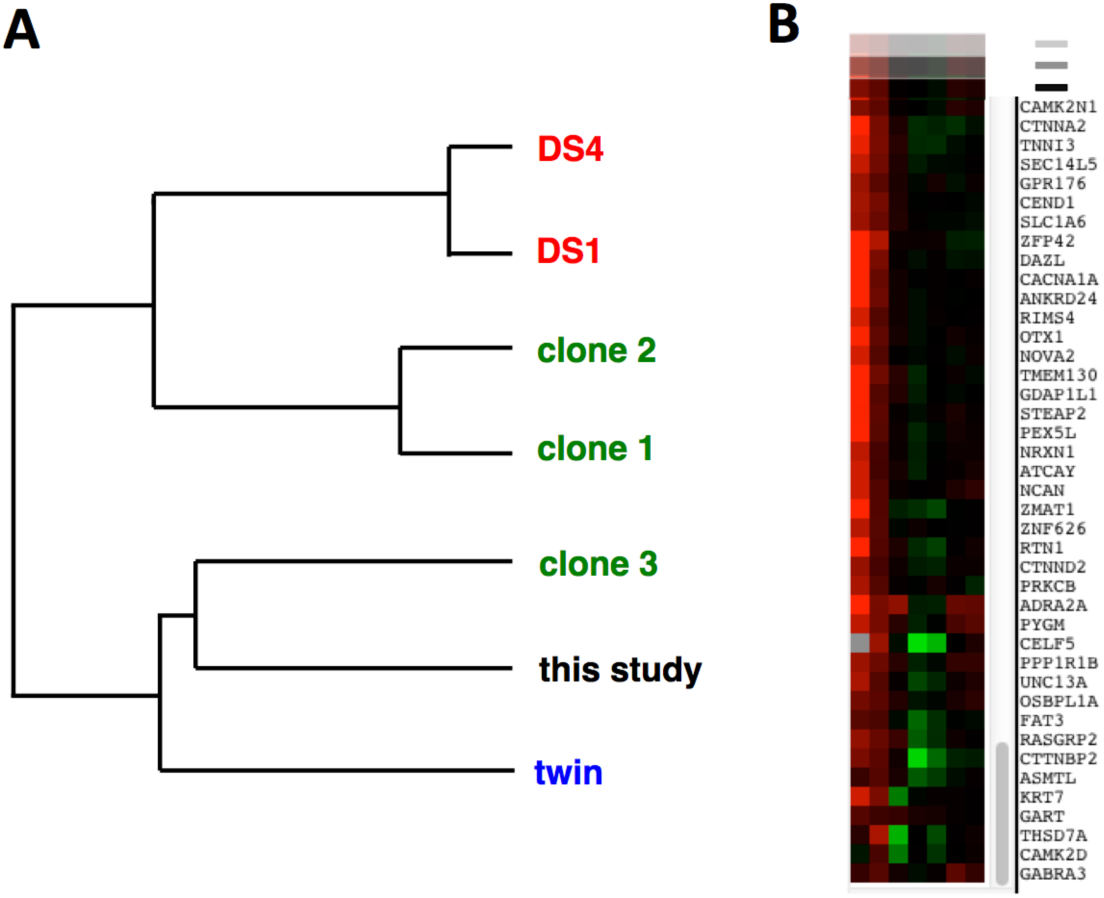
Cluster analysis of comparable studies using isogenic disomic/trisomic cell pairs. **A.** Dendrogram representation of relatedness of datasets listed in Table 2. DS1 and DS4, from Weick et al.; clones 1,2,3 from Jiang et al; “twin” from Letourneau et al. **B.** Section of heat map display showing expression ratios for individual genes calculated for datasets shown in (**A**.). Note overall similarity between “twin” data (leftmost column) and the data from this study (next column).

## Discussion

### Transcript accumulation differences between cells trisomic or disomic for chromosome 21

An inherent limitation of human transcriptome studies is accounting for the differing genetic background of individuals. In the context of Down syndrome, this limitation has hindered determining the degree to which trisomy of chromosome 21 alters expression of genes on this chromosome and in the genome as a whole. However, recent technical advances have allowed the derivation of iPSCs trisomic or disomic for chromosome 21 from the same individual. We have used RNA-seq to characterize in depth the transcriptome of one such trisomic/disomic iPSC pair, originally generated by Li and colleagues (Li et al, 2012). We have also differentiated these iPSCs into cortical neuronal cultures to investigate how trisomy of chromosome 21 impacts the transcriptome in differentiated neurons. Our results indicate that an increase in chromosome 21 dosage does generally lead to increased transcript accumulation for chromosome 21 genes. However, not all chromosome 21 genes follow this pattern, and which genes show increased expression depends on the cell type assayed (iPSCs vs. differentiated cortical neurons). This observation may not be surprising, given how regulatory loops, compensatory changes, and cell-specific transcriptional programs may modulate or override the dosage effect on any given chromosome 21 gene. In this regard, it has recently been reported (Sullivan et al, 2016) that the interferon response is up-regulated in trisomy 21 fibroblasts and lymphoblast lines, likely due to increased expression in these cells of the four interferon receptors (IFNAR1, IFNAR2, IFNGR2, and IL10RB) located on chromosome 21. We find that these receptors also have increased transcript accumulation in the trisomic iPSCs we analyzed, but this is not apparent in the derived neuronal cultures.

As observed in other studies, the majority of significant transcriptome changes we identified in both iPSCs and derived neuronal cultures occur in non-chromosome 21 genes. Gene ontology analysis of genes with transcript abundance changes in trisomic iPSCs unexpectedly revealed increases in transcripts from neuronal-associated genes and decreases in transcripts from genes involved in cell adhesion function and germ layer/mesoderm development. These differences did not result from visible spontaneous differentiation in the trisomic iPSCs, which were monitored daily. We note, however, that cells in iPSC cultures are not homogeneous (see Figure 2), and one interpretation of our results is that the trisomic iPSC cultures contain higher proportions of stem cells biased towards neuronal development.

One significant consideration in comparing the trisomic and disomic iPSCs used in this study is that the disomic clones have gone through two cycles of selection: first with G418 to select for the insertion of a dual selection cassette into exon 3 of the *APP* gene, and then with gancyclovir to select for loss of the chromosome 21 bearing the inserted selection cassette. It is possible that these rounds of selection resulted in epigenetic (or, less likely, genetic) changes that added to the transcriptional differences between the trisomic and disomic iPSCs. In fact, we were unable to efficiently differentiate into cortical neuronal cultures any of the 4 disomic clones that had gone through these selection procedures. As the differentiation protocol we used has been employed by other research groups to produce cortical neuron cultures from iPSCs with normal karyotypes, we suspect that the derivation of disomic clones may have resulted in a subtle intrinsic change in the iPSC state. Fortunately, we were able to derive a spontaneous disomic line from the C3 trisomic iPSC clone, which did readily differentiate into cortical neurons. The C3 disomic iPSCs (C3-D21) were therefore used to generate the disomic cortical neuronal cultures. Importantly, C3-D21 disomic iPSCs expressed less MAP2 than trisomic iPSCs (Figure 3C), and we still observed increased expression of neuronal-associated genes in the C2 trisomic neuronal cultures compared to the disomic neuronal cultures derived from C3-D21. These observations argue that our finding that C2 trisomic cells have a more neuronal character than disomic cells from the same individual cannot be attributed to an artifact of the genetically-selected disomic clones.

The neuronal differentiation protocol we used (Espuny-Camacho et al, 2013) has been reported to result in the production of layer V and VI cortical neurons after 40 days in culture, which we confirmed by immunofluorescence microscopy using antibodies against layer V and VI - specific transcription factors CTIP2 and TBR1, as well as antibodies to glutamate receptor GRIK2 and glutamate transporter vGLUT1 (Supplementary Figure 1). While the neuronal cultures derived from the trisomic and disomic iPSCs were not readily distinguishable using any of these markers, there were significant transcriptome differences. Of particular note was increased transcript accumulation for synaptic proteins, including GABA (GABRG2, GABRA1, GABRB2, GABBR2) and glutamate (GRIN2B, GRIA1) receptors in the trisomic iPSC-derived neuronal cultures. We note that increased GABAnergic signaling occurs in the Ts65Dn mouse DS model (Kleschevnikov et al, 2012a), and counteracting GABAnergic inhibition in these mice reverses behavioral deficits (Kleschevnikov et al, 2012b).

### Transcriptome characteristics not affected by chromosome 21 trisomy

The increased chromosome 21 gene dosage in trisomic cells could conceivably affect global RNA metabolism, as chromosome 21 encodes the RNA editing gene ADARB1 and multiple splicing factors (including U2AF1, SON and SCAF4). Global chromatin effects of chromosome 21 trisomy were also proposed by Letourneau et al (2014), who identified gene expression dysregulation domains (GEDDs) in trisomic iPSCs and fibroblasts. We have analyzed our transcriptome data for trisomy-dependent changes in adenosine-to-inosine RNA editing, alternative splicing, repetitive element expression, and chromosomal domains of altered expression. Examining more than 10,000 annotated RNA editing sites (from the DARNED database), we did not find any evidence for global changes in RNA editing in trisomic iPSC-derived neuronal cultures. This analysis does not preclude changes in site-specific editing, which would require significantly deeper sequencing depth to assay accurately. Nevertheless, our data indicate that global editing levels do not correlate with ADARB1 transcript levels, likely due to the multiple factors that control the accumulation of edited transcripts. Our data also did not reproduce the gene expression domains identified by Letourneau et al (2014), and we note that an independent analysis of the Letourneau data has questioned the existence of GEDDs (Do et al., 2015).

### Altered splicing in trisomic cells

We did observe trisomy-dependent splicing changes in both the iPSCs and neuronal cultures. Our identification of splicing alterations in trisomic cells is not completely novel, as splicing differences in a selected set of genes has been observed in fetal DS tissue (Toiber et al, 2010). RNA-seq has also been used previously to support altered splicing in DS endothelial progenitor cells, although no confirmatory studies were done (Costa et al, 2011). The large majority of splicing changes we identified did not occur in genes located on chromosome 21,suggesting that they were not a direct effect of gene dosage (i.e., higher gene dosage leading to competition for splicing factors and subsequent altered splicing). More likely is the possibility that altered expression or altered activity of splicing factors caused the splicing dysregulation observed in trisomic cells. We did observe increased transcript accumulation for three known or suspected splicing factors (U2AF1, SON, and SCAF4) located on chromosome 21 in the trisomic iPSC and/or neuronal cultures. Alternatively, increased expression of chromosome 21 gene DYRK1A, a kinase known to regulate splicing factors (Qian 2011), could contribute to the splicing changes we identified. Over-expression of DYRK1A in mice has been reported to mimic Down syndrome splicing aberrations (Tobier et al, 2010). Direct demonstration of a role of DYRK1A or specific splicing factors in the trisomy-dependent splicing changes we have observed will require additional studies in which the gene dosage or expression levels of these genes are normalized in trisomic backgrounds.

### Altered HERVH expression in trisomic iPSCs

Examination of the accumulation of repetitive element transcripts revealed a highly significant decrease in transcripts from the endogenous retrovirus HERVH in the trisomic iPSCs. HERVH transcripts have been specifically identified as a pluripotency marker (Santoni et al, 2012), and disruption of HERVH transcription compromises self-renewal in human embryonic stem cells (Wang et al, 2014). The reduced HERVH expression we observe in trisomic iPSCs is consistent with these cells having an altered pluripotent state in comparison to the disomic iPSCs.

### Comparison with other datasets

As described above, consideration of the possible effects of the selection procedures used to generate the disomic derivative clones suggests a possible technical reason why we have observed transcriptional differences not reported by other investigators. However, cluster analysis of our dataset with other comparable studies suggests that our results are not anomalous, but may reflect the range of states assumed by human stem cells trisomic for chromosome 21. The dendrogram shown in figure 6A could be interpreted as showing that there are two classes of trisomic stem cells: one class with a more neuronal character (cells from this study, the twin study of Letourneau et al, and clone 3 from the Jiang study), and one class with a more mesodermal character (clones 1 and 2 from the Jiang study, and both trisomic lines from the Wieck et al study). This is likely an oversimplification; more extensive data might readily reveal a spectrum of trisomic stem cell fates. However, this analysis does suggest that there is unlikely to be a single transcriptome signature for trisomic stem cells, and the observed transcriptome variation may not simply be a result of varying experimental conditions. This latter point is supported by the observation that clone 3 of the Jiang et al study clustered more closely with our data set than with the other XIST clones. The Jiang study was particularly well controlled, as the iPSCs were grown under the same conditions (using mouse embryonic fibroblasts as feeder cells) with the same length of XIST-based knockdown of chromosome 21 expression (3 weeks). Nevertheless, clone 3 trisomic iPSCs appeared more similar to ours, which were grown in feeder-free media.

### Chromosome 21 trisomy and stem cell pluripotency

Why might trisomic iPSCs show a range of transcriptome signatures? Given our observation that the trisomic iPSCs we characterized had enhanced accumulation of transcripts associated with differentiated neurons, and reduced expression of the HERVH pluripotency marker, we speculate that trisomic iPSCs may have an inherent inability to maintain a full pluripotent state. Following this assumption, trisomic iPSCs would have a more constrained differentiation capacity than corresponding disomic iPSCs, which could explain their reported defects in neuronal differentiation (Jiang et al, 2013). On the other hand, in the absence of full pluripotency, trisomic iPSCs might alternatively be biased towards ectodermal fates (e.g., the trisomic iPSCs we have characterized). The specific developmental bias of a given trisomic iPSC might depend on a variety of factors: genetic background, initial culture conditions, pluripotency factor expression, or stochastic variation. If the gene dosage burden imposed by chromosome 21 trisomy generally interferes with the maintenance of stem cell fate, this might account for some Down syndrome clinical phenotypes, including the hematopoietic developmental defects observed in DS (Roberts and Izraeli, 2014). In this regard, it has been reported that the Ts65Dn mice have defects in the self-renewal of hematopoietic stem cells as well as in the expansion of mammary epithelial cells, neural progenitors and fibroblasts, specifically due to the triplication of Usp16 (Adorno et al, 2013).

---

**Supplemental Figure 1.**
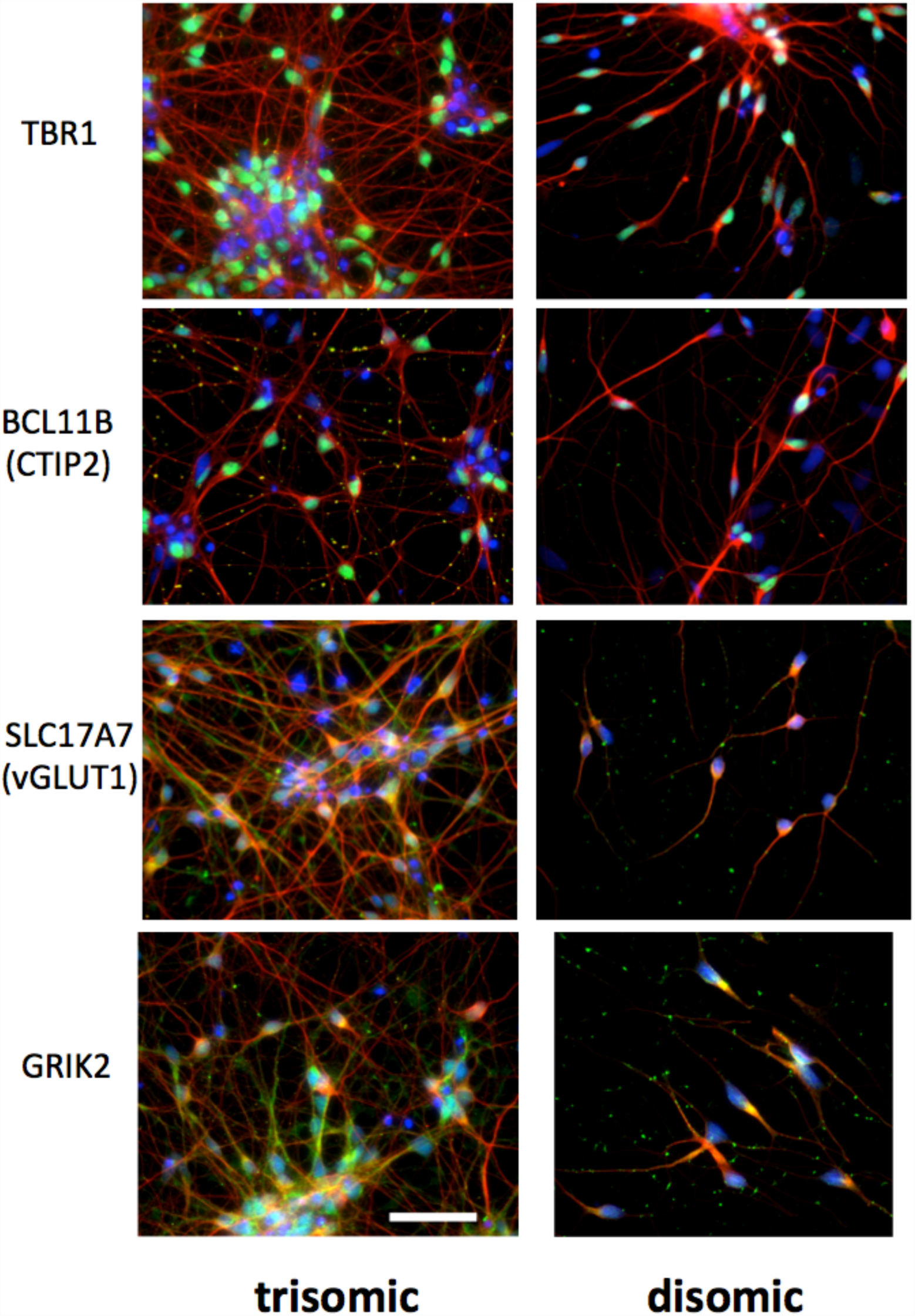
Confirmation of differentiation of iPSCs into cortical neuronal cultures. Images taken from clones C2 (trisomic) and C3 -D21 (disomic) 40 days after initiation of the differentiation protocol. Fixed cells on chamber slides were probed with antibodies against the marker proteins listed in the left column. Neuronal marker Beta III tubulin is red; DAPI is blue; and Ctip2, TBR1, SLC17A7 (vGLUT1) and GRIK2 are green. Size bar = 20 microns.

**Supplementary Figure 2.**
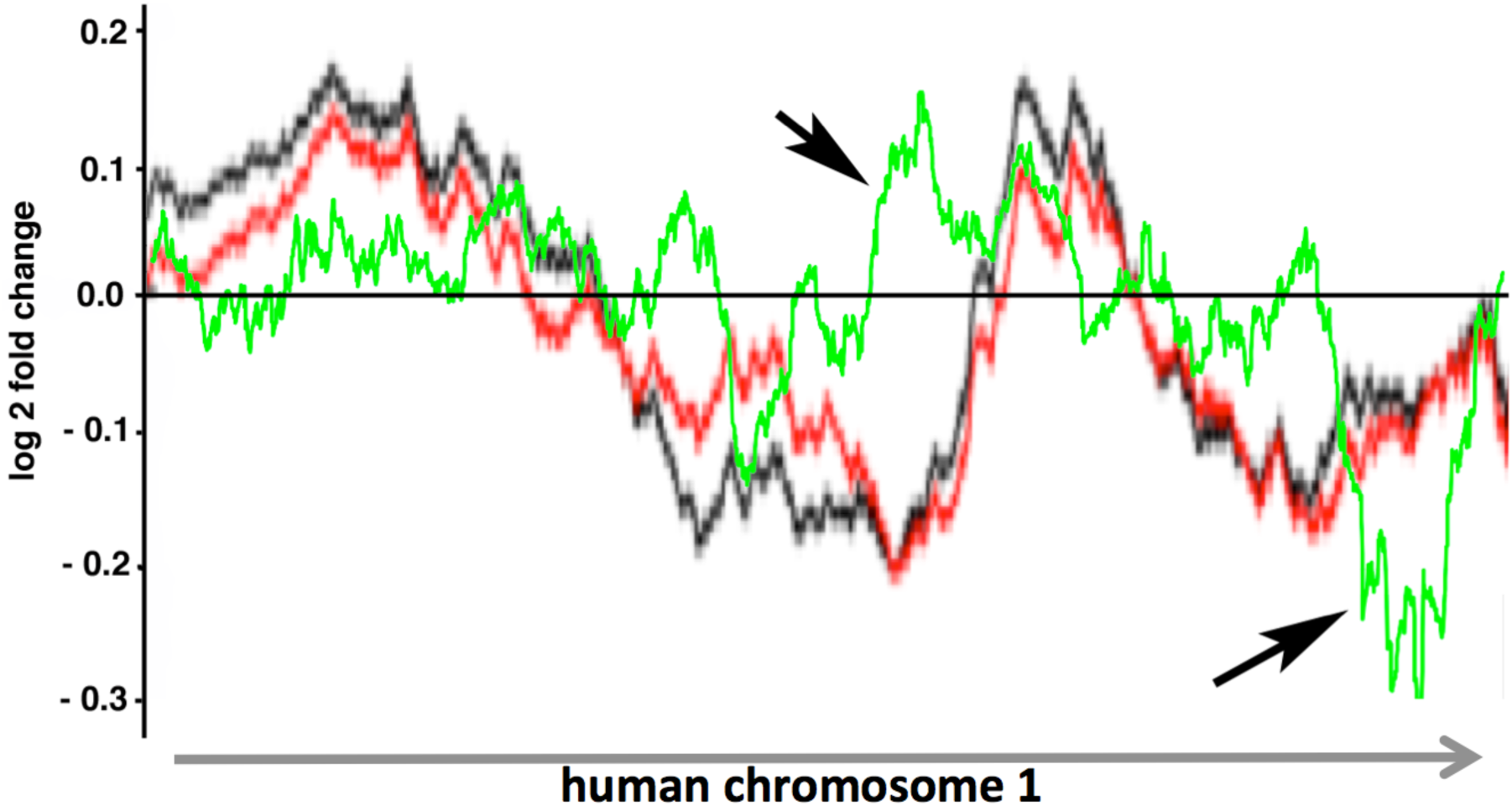
Comparison of putative trisomy-dependent gene expression domains identified on chromosome 1 by Letourneau et al (2014) and this study. Lines show regions of increased or decreased gene expression in cells trisomic for chromosome 21 as calculated using the smoothing algorithm employed by Letourneau et al. The red (iPSCs) and black (fibroblast) traces are taken from the Letourneau study; the green trace is from the iPSCs used in this study. Note that there are significantly different expression domains calculated from the different studies (arrows).

